# Chimpanzee (*Pan troglodytes verus*) density and environmental gradients at their biogeographical range edge

**DOI:** 10.1101/2020.07.14.202499

**Authors:** Erin G. Wessling, Paula Dieguez, Manuel Llana, Liliana Pacheco, Jill D. Pruetz, Hjalmar S. Kühl

**Affiliations:** Max Planck Institute for Evolutionary Anthropology, Deutscher Platz 6, 04103 Leipzig, Germany; Harvard University, 11 Divinity Ave, Cambridge, MA, USA; Instituto Jane Goodall España, Station Biologique Fouta Djalon, Dindéfélo, Région de Kédougou, Senegal; Texas State University, San Marcos, Texas, USA

**Keywords:** niche, habitat, food availability, ecography, thermoregulation, savanna-mosaic, species range limit

## Abstract

Identifying ecological gradients at the range edge of a species is an essential step in revealing the underlying mechanisms and constraints that limit the species’ geographic range. We aimed to describe the patterns of variation in chimpanzee (*Pan troglodytes verus*) density and habitat characteristics perpendicular to the northern edge of their range and to investigate potential environmental mechanisms underlying chimpanzee distribution in a savanna-mosaic habitat. We estimated chimpanzee densities at six sites forming a 126 km latitudinal gradient at the biogeographical range edge of the western chimpanzee in the savanna-mosaic habitats of southeastern Senegal. To accompany these data, we used systematically placed vegetation plots to characterize the habitats at each site for habitat heterogeneity, tree density and size, floral assemblages, among other variables. We found that both biotic and abiotic factors are potential determinants of the chimpanzee range limit in this ecoregion. Specifically, chimpanzee-occupied landscapes at the limit had smaller available floral assemblages, less habitat heterogeneity, and contained fewer closed canopy habitats in which chimpanzees could seek refuge from high temperatures than landscapes farther from the range limit. This pattern was accompanied by a decline in chimpanzee density with increasing proximity to the range limit. Our results provide several indications of the potential limits of food species diversity, thermal refuge, and water availability to the chimpanzee niche and the implications of these limits for chimpanzee biogeography, especially in the face of climate change predictions, as well as to species distributional modeling more generally.

## INTRODUCTION

Abundant-center niche theory predicts that habitat suitability is highest at the center of a species’ range and decreases towards the outer boundaries of where that species is found (Hutchinson, 1961; Brown 1984; Brown et al.1995; Holt, 2009). As habitat suitability decreases, species density likewise is expected to decrease, so that densities should be lowest at the range edge (Kawecki 2008; Sexton et al. 2009). Patterns of species density distributions can vary considerably depending on the limiting factors that dictate them. Accordingly, species densities may decline gradually across the range towards the edges or may remain stable across the range until they drop off at the very limits of niche tolerance (Brown 1984; Brown et al.1995). Additionally, many species’ ranges extend outside suitable habitat into marginal habitats. Marginal habitats represent a transition zone or ‘bleed-over’ of individuals from suitable habitats into unsuitable habitats and allow for little margin of variation of environmental factors before they become uninhabitable (Kawecki 2008). As such, these marginal habitats may become population sinks, solely supported by immigration from more suitable habitats (Pulliam 1988; Marshall 2009; Smith et al. 2011). The gradient of habitability at the range edge may thus represent an important natural scenario in which to investigate environmental drivers of biogeographic limitations.

Drivers of biogeographic limitations are typically investigated in small-bodied organisms (e.g., Chown and Gaston 1999; Hargreaves et al. 2014; Cahill et al. 2014), likely because investigating range constraints in large-bodied, long lived species, like many primates, is challenging. Despite the challenges, as the underlying processes of range biogeography may differ between short- versus long-lived species due to differences in their life history strategies, investigating range constraints of long-lived species is essential for a comprehensive understanding of these processes. For example, the importance of fallback foods in defining habitat carrying capacity for long-lived species may be unique because theoretically they prioritize survival more than short-lived species, who typically prioritize high reproductive rates (Marshall et al. 2009a).

For example, many frugivores are able to track fruiting patterns at small spatial and temporal scales, and fruit availability can be a driver of frugivorous species’ abundance (White 1978). These patterns have been investigated in a number of frugivorous bird species (e.g., Rey 1995; Restrepo, Gomez, and Heredia 1999; Moegenburg and Levey 2003; Seoane et al. 2006), various ape (e.g., Vogel et al. 2015; Pennec et al. 2016; Marshall and Leighton 2006; Marshall et al. 2014), and various primate species (e.g., Kinnaird and O’Brien 2005; Milton et al. 2005; Rovero and Struhsaker 2007). However, climatic constraints to species biogeography are the most common of range constraints and are thought to be specifically limiting to a number of taxa, including primates (Korstjens et al. 2010; Stone et al. 2013). For example, areas rich in food resources have even been ignored by baboons if they were too far from water sources (*Papio cynocephalus*: Altmann 1974; *Papio ursinus*: Hamilton et al. 1976), suggesting food availability may become irrelevant to species otherwise constrained climatically.

The relative importance of environmental components may shift from within the range towards its limits, regardless of habitat suitability. The northern limit of the western chimpanzee range therefore presents a unique opportunity to investigate environmental correlates of distribution in a large-bodied long-lived primate. While geographic barriers (e.g., the ocean, the Congo River, disturbance in the Dahomey gap) limit many edges of the chimpanzee biogeographic range and remaining natural edges are rare, the northern limit of this subspecies’ range is otherwise unhindered by immediate geographic barriers. Limitations to chimpanzee distribution at this range edge have been linked to thermoregulatory limitations (McGrew et al. 1981), decreased floral richness and diversity (Kortlandt 1983), time constraints (Korstjens et al. 2010), and water scarcity (Lindshield 2014), although none have been extensively investigated.

A number of large-scale chimpanzee ecological niche models (ENMs), or distribution models, have offered insights into the factors that influence chimpanzee distribution and site occupancy, such as distance to roads or rivers, forest coverage, climatic and human influences (Junker et al. 2012; Sesink Clee et al. 2015; Foerster et al. 2016; Jantz et al. 2016; Abwe et al. 2019; Heinicke et al. 2019a; Barratt et al. 2020), although these models typically evaluate characteristics of the chimpanzee niche at large. Larger-scale analyses depend on data obtained from remote sensing (e.g., percent forest cover, climate averages, human population indices, land cover classifications, distance to roads and rivers) as proxies for smaller-scale metrics like habitat heterogeneity or food species assemblages. Small-scale metrics can offer insights into more direct, proximate drivers of chimpanzee distribution and niche suitability but are rarely included in these models as they are either unavailable or not easily inferable from the methods used for larger-scale analyses (but see Foerster et al. 2016). For example, chimpanzees are unlikely to evaluate habitat for suitability based on percent forest coverage in the landscape (e.g., McGrew et al. 1988; Junker et al. 2012; Heinicke et al. 2019a), but percent forest cover may be a proxy of potential shade and food resources available which would be directly relevant to chimpanzee daily life and space use. In this way, smaller-scale studies can offer a more direct understanding of proximate mechanisms underlying species occupancy variation than large-scale studies, including for chimpanzees.

Finer-scale correlates of chimpanzee density variation have been extensively investigated in the forested habitats of East and Central Africa (Balcomb et al. 2000; Potts et al. 2009; Bortolamiol et al. 2014; Foerster et al. 2016; Potts and Lwanga 2013; Potts et al. 2015; Nguelet et al. 2016). As chimpanzees prefer ripe fruit over other food types (Wrangham 1977; Conklin-Brittain et al. 1998), evaluated predictors of chimpanzee densities often center on the availability of this food. Variation in the availability of fruit resources has been linked to various aspects of chimpanzee social life (e.g., Chapman et al. 1995; Boesch 1996; Murray et al. 2006; Wittiger & Boesch 2013; Samuni et al. 2018), physiology (e.g., Wessling et al. 2018a,b; Emery Thompson et al. 2009, 2010) and subsequent reproductive output (e.g., Pusey et al. 1997; Emery Thompson et al. 2007). Therefore, it follows that variation in these resources should impact chimpanzee abundances.

Some authors posit that tree densities likewise play a role in shaping this range limit in the sense that closed canopy habitats (e.g., gallery forests) within these landscapes offer refuge from the hot and difficult climate conditions (McGrew et al. 1981). In contrast, that chimpanzees rely on diverse plant assemblages to support their diverse diet, and that an ultimate characteristic of the chimpanzee range limit in savanna-mosaics is argued to be strictly related to plant species richness (Kortlandt 1983). Further, the extent to which fruit species density dictates chimpanzee distribution in the savanna-mosaics of West Africa is unclear as factors like thermoregulation and dehydration play stronger seasonal roles than energetic constraints at the individual level in these habitats (Wessling et al. 2018b).

Savanna-mosaic habitats are predicted to harbor lower tree densities overall compared to forested habitats (Crowther et al. 2015), but a comparison of isotopic evidence perpendicular to the range limit (i.e., with decreasing distance to the range limit) failed to identify patterns of isotopic variation that would suggest that chimpanzees at the range limit experience higher degrees of nutritional scarcity or starvation (Wessling et al. 2019). Instead, the authors suggested that chimpanzees at the range limit compensate for potential nutritional scarcity by using fallback foods that are high in δ^13^C, such as flowers, domestic cultivars, or grasses. However, the prediction that chimpanzees at the range limit experience higher degrees of nutritional scarcity than those further from the range limit is predicated on the assumption that habitat suitability decreases with proximity to the limit. This assumption remains untested, and the authors’ isotope results may indicate that these sites (Fongoli, Kanoumering, Makhana, Kayan, and Hérémakhono) did not vary in their ability to support chimpanzees (Wessling et al. 2019).

We set out to investigate the nature of the chimpanzee biogeographical range edge in southeastern Senegal. Specifically, we aim to describe patterns of variation in chimpanzee density and habitat characteristics perpendicular to their biogeographical range limit, and to investigate potential environmental candidates for the structure of the chimpanzee range edge in this savanna-mosaic landscape. We define the range limit as the last biogeographical point at which a species (in this case, chimpanzee) can be found, and a range edge as the region near to the range limit at which species densities decline. This decline may occur over the large or small scale and may occur across a single or multi-dimensional niche space (Sexton et al. 2009). We hypothesize that chimpanzee habitat suitability decreases with increasing latitude (and therefore proximity to the presumed range limit), and therefore in turn chimpanzee densities likewise decrease. As habitat suitability can be characterized in a number of ways, we specifically predict that (1) tree density, (2) tree size (i.e., DBH), (3) number of available food species, (4) proportion of trees within preferred food categories, and (5) available refuge habitats (i.e., closed canopy habitats) decrease with increasing latitude (i.e., proximity to the distributional limit).

As common determinants of species’ range limits, abiotic factors such as temperature or rainfall may be stronger determinants of the range than biotic metrics (which are ultimately also shaped by abiotic conditions). Therefore, we consider the alternative hypothesis that abiotic factors directly determine the chimpanzee range limit in savanna-mosaic habitats (McGrew et al. 1981), with the prediction that temperature increases and rainfall decreases towards the range limit, as implied by long-term national-level climatic models (Fall 2006). These two hypotheses (abiotic vs. biotic determinants) are not mutually exclusive but are likely to contribute collectively as a suite of habitat characteristics to the definition of the range limit. We cannot address the relative strength of support for each hypothesis in this study, but rather investigate the available support for each. The advantage of our approach is that we investigate these processes within a single ecoregion (i.e., savanna-mosaic woodland), and therefore are unburdened by other potentially confounding variables attendant to large-scale ENMs.

In this investigation we must assume that the limit is determined to some degree by naturally occurring bottom-up processes (e.g., food availability) and not solely by top-down processes (e.g., predator abundance or anthropogenic disturbance). Anthropogenic disturbance has had historical effects in the study region over decades (e.g., Mbow et al. 2000; Tappan et al. 2004) and presumably continues to play a role in shaping biotic landscape metrics. However, as hunting chimpanzees is a regional taboo (Heinicke et al. 2019a) and therefore anthropogenic influence in chimpanzees is predominantly indirect (through its effects upon the environment), our study investigates the bottom-up processes (including potential anthropogenic factors) dictating chimpanzee range limits.

## METHODS

We collected data at six sites located along a latitudinal gradient in southeast Senegal (Figure 1; Table 1). This region of Senegal is located in the Shield ecoregion (Tappan et al. 2004) and is generally described as a highly seasonal savanna-woodland mosaic comprising gallery forest, woodland, and grassland. It is considered particularly extreme in comparison to other habitats where chimpanzees are studied due to the extensive dry season and comparatively hot temperatures (Pruetz & Bertolani 2009; Wessling et al., 2018a). The six sites in this study include the habituated research group of the Fongoli Savana Chimpanzee Project (FSCP), two unhabituated chimpanzee research sites (Kayan, and the RNCD: Réserve Naturelle Communautaire de Dindéfélo, hereafter Dindefelo) of the Pan African Programme (PanAf: http://panafrican.eva.mpg.de), and three additional unhabituated chimpanzee sites (Kanoumering, Makhana, Hérémakhono). Researchers have monitored the chimpanzees at Dindefelo since 2009, prior to the adoption of the PanAf protocol in 2016. Our discovery of chimpanzees at Hérémakhono led to amendments of the IUCN range limits for the western chimpanzee, as chimpanzee presence had not been previously verified north of the Parc National de Niokolo Koba (Figure 1; Humle et al. 2008; Humle et al. 2016). Based on data collected via interviews and reconnaissance surveying at eight sites north of the Hérémakhono site, data suggest that Hérémakhono represents the very northern-most vestige of chimpanzee distribution or is very near to it (Wessling et al. 2019). These results match findings by Lindshield et al. (2014) who surveyed an additional five sites north of Hérémakhono to confirm chimpanzee absence. While it is possible that chimpanzees continue to range north of Hérémakhono, these survey campaigns were specifically directed at range limit discovery and failed to detect chimpanzee presence in any locality to the north of our study area, supporting the idea that Hérémakhono likely represents or is near to the northern-most location in which chimpanzees range.

**Figure 1.**
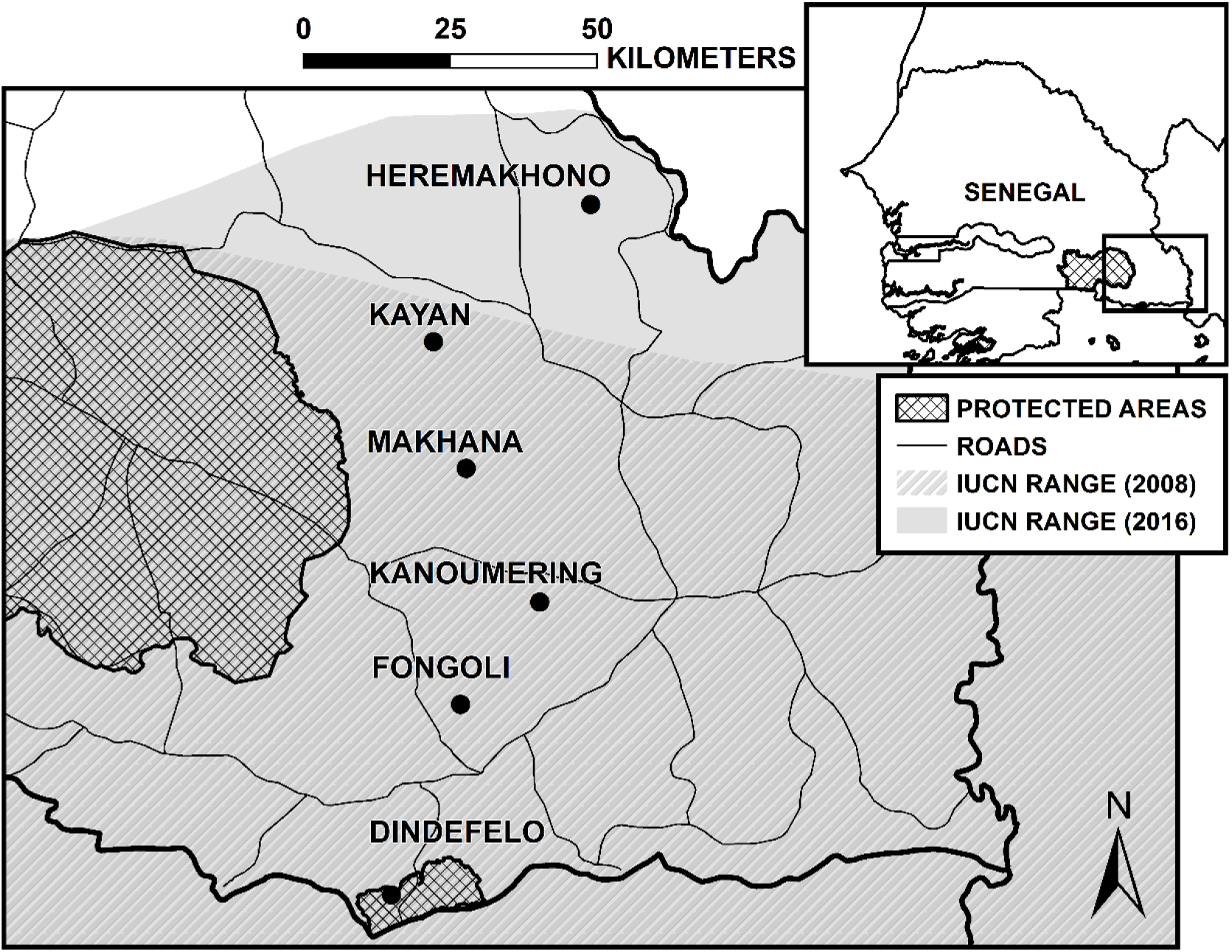
Map of the study area, with six study sites, relative to protected areas, and the current and former IUCN *P.troglodytes verus* range limit (Humle et al. 2008, 2016).

**Table 1.** Estimates of chimpanzee (*Pan troglodytes verus*) density, survey lengths, and climate characteristics for each of the six sites in this study (2012-2017), ordered from north (top) to south (bottom). Significant (p<0.05) latitudinal patterns are indicated in underscored bold text, based on Spearman correlations between latitude and the variable of interest.

| Site | Longitude / Latitude | Number of grid cells | Total distance surveyed (survey length; km) | Total number of nests encountered on transects (on recces) | Chimpanzee density (± SE; indiv. / $\text{km}^2$) | Mean daily temperature ($^{\circ}$ C) | Mean daily maximum temperature ($^{\circ}$ C) | Mean annual rainfall (measured rainfall; mm) |
| --- | --- | --- | --- | --- | --- | --- | --- | --- |
| Hérémakhono | 823665 / 1486000** | 20 | 40.0 (20.0) | 0 (21) | <u>**0.00**</u> | 30 | 37 | <u>**849**</u> |
| Kayan | 797836 / 1462391 | 74 | 94.0 (31.3) | <u>66 (410)</u> | <u>**0.05 (0.04 – 0.05)**</u> | 28 | 37 | <u>**906 (1109)**</u> |
| Makhana | 803500 / 1441000 | 20 | 60.0 (20.0) | <u>66 (350)</u> | <u>**0.07 (0.07 – 0.08)**</u> | 30 | 39 | <u>**966**</u> |
| **Kanoumering** | 816000 /<br>1418500 | 20 | 60.0<br>(20.0) | <u>81 (481)</u> | <u>0.09 (0.08 – 0.10)</u> | 30 | 35 | <u>1029</u> |
| **Fongoli** | 803000 /<br>1401000 | 20 | 20.0<br>(20.0) | <u>89 (*)</u> | <u>0.29 (0.26 – 0.33)</u> | 29 | 37 | <u>1086</u><br><u>(1016)</u> |
| **Dindefelo** | 791841 /<br>1368583 | 79 | 73.8<br>(36.9) | <u>147 (*)</u> | <u>0.13 (0.12 – 0.15)</u> | 29 | 34 | <u>1220</u><br><u>(1243)</u> |
| ***p - value*** | - | - | - | - | <u>0.017</u> | 0.64 | 0.23 | <u><0.01</u> |
| ***rho*** | - | - | - | - | <u>-0.94</u> | 0.25 | 0.58 | <u>-1.00</u> |

\* Recces were not surveyed in these permanent research sites
\*\* Coordinates approximated as this study site overlaps two UTM zones (28P and 29P)

Two sites (Kayan and Makhana) are relatively close to Parc National de Niokolo Koba, and no villages or roads exist between the study areas and the boundaries of the Park, and Dindefelo is a formally-recognized community reserve. Hérémakhono and Fongoli, however, are each located near to one or more villages and include minor degrees (<5% land cover; Wessling unpublished data; Bogart & Pruetz 2011) of anthropogenic habitats within the landscapes (e.g., agricultural fields). Kanoumering, while comparatively remote, had minor degrees of human foot traffic and disturbance for gold mining exploration at the time of our research, but appeared otherwise undisturbed by land conversion. In total, these six sites represent approximately the northern 126km of the range edge in West Africa. We noted impacts of bush fires and felling for livestock at all six sites. Other indications of human activity included vehicle roads for former (Makhana) and ongoing (Fongoli) gold exploration, artisanal logging (Hérémakhono, Fongoli, Dindefelo) and gold mining (Fongoli, Hérémakhono), and palm wine collection (Kanoumering). These observations indicate that all six sites suffer from some degree of anthropogenic disturbance, although we do not quantify these patterns here.

### Chimpanzee Density and Habitat Data Collection

In the cases of previously un-surveyed sites (Kanoumering, Makhana, Kayan, Hérémakhono), initial reconnaissance (recce) surveys were conducted to identify contiguous areas of chimpanzee presence at each site. A 1 km by 1 km contiguous grid system was overlaid at each site to contain locations in which nests were discovered during recce surveys until at least 20 grid cells had been placed. We chose 20 cells as a minimum to maintain consistency with the PanAf protocol (http://panafrican.eva.mpg.de/) minimum. For the two pre-existing sites (Fongoli and Dindefelo), a minimum number of 20 grid cells were overlaid on known chimpanzee community home ranges. Continuous presence at the two PanAf sites (Kayan and Dindefelo) allowed us to extend the grid system over a greater extent of chimpanzee home range use (up to 79 km^2^). Despite site differences in survey area, all subsequent measures described in this manuscript account for differences in research area. Data at all sites were collected over one annual cycle. We used chimpanzee nests, commonly used as signs of chimpanzee presence (Kühl 2008), to estimate chimpanzee densities at each site. Along the north to south mid-point of each grid cell, straight line transects were walked to estimate chimpanzee abundance using distance sampling (Buckland et al. 2001; Kühl 2008), and the perpendicular distance from the transect to each discovered nest was recorded. In Kayan and Dindefelo where the research area was extensive, transects were walked in alternating grid cell lines (i.e., 2 km longitudinal distance between continuous transects). Each set of transects were subsequently resurveyed 1-2 times at intervals between two and eight months except for Fongoli which was sampled only once due to time constraints (Table 1). Line transects were predominately surveyed during the dry season (October – April) under conditions of good visibility, per standard surveying protocols (Kühl 2008). However, sites which were surveyed three times (Kanoumering, Makhana, and Kayan) were surveyed once during the wet season (June – September) to maintain even temporal distribution; wet season perpendicular distances did not markedly differ from dry season rounds at these sites although nest discoverability was considerably lower (mean of 41% of dry season counts).

We calculated chimpanzee densities (D) using the equation: D = N/(2 ∗ L ∗ ESW ∗ p ∗ r ∗ t), where N is number of nests discovered within the truncation distance, L is the length of the transect, ESW is the effective strip width, p is the proportion of nest builders, r is the nest production rate, and t is the nest decay rate (Buckland et al. 2001; Kühl 2008). We calculated ESW based on perpendicular distances from the transect at Makhana, Kanoumering, and Fongoli using the ‘Distance’ package (Miller 2017; Thomas et al. 2010) in the statistical software R (version 3.6.1; R Core Team 2019). We could not include Dindefelo, Kayan, or Hérémakhono in our calculation of ESW as perpendicular distances had not been recorded at these sites, and therefore based density estimations for all sites based on a single pooled ESW. This method assumes that nest discoverability remains constant across the savanna-mosaic ecoregion (i.e., areas where ecosystems are generally similar; Buckland et al. 2001) and averages out potential stochastic (i.e., random) influences that may have arisen with smaller, locally scaled site-based datasets. We used 63.22 m as the truncation distance when calculating ESW, as this was the maximum perpendicular distance observed once we excluded a single extreme outlier (183 m) from the dataset. To complete the equation, we used a nest production rate of 1.142 nests per individual (Kouakou et al. 2009), 0.83 as the proportion of nest builders (Plumptre and Cox 2006), and a nest decay rate of 243 days per nest based on data collected in Dindefelo (Heinicke et al. 2019a). To ensure that wet season surveying did not impact our evaluation of latitudinal patterns on chimpanzee density, we also estimated chimpanzee densities at sites which had been surveyed during the wet season, with wet season surveys excluded. Although densities at these sites were slightly higher based only on dry season surveys, this modification had no effect on latitudinal patterns in chimpanzee densities and we therefore report results from our complete surveys only.

In addition to straight line transects for chimpanzee density calculations, we also collected environmental data for each site using vegetation plots centered along these transects. Diameter at Breast Height (DBH), location, and species identification data were collected for trees and lianas 10 cm or larger DBH in 20 by 20 m plots spaced at 100 m meter intervals. Due to the extent of the grid systems at the PanAf sites, vegetation plots were instead placed either at one corner of each grid cell and at the end of transects (Kayan) or at 200 m intervals (Dindefelo). Data collection in logistically unfeasible (e.g., steep rock faces) vegetation plots was abandoned (n=14). We mistakenly missed sampling a single plot at Makhana and additionally sampled at Hérémakhono in error. We did not sample vegetation plots at Fongoli due to time constraints. Data collected from a 3.4 km by 20 m phenology transect placed randomly within the Fongoli home range (traversing but not parallel to the transect for nest surveying) can nonetheless offer an estimate of DBH and tree genera composition. Due to difficulty identifying species within specific genera (e.g., *Acacia, Ficus*), all subsequent analyses operate at the level of the genus. We calculated basal area (BA) as the sum of the basal area (area = (0.5 ∗ DBH)^2^ ∗ π) of all trees in the site, divided by area of vegetation plots surveyed.

Lastly, we collected year-round data on daily temperature at each site using a min-max hygrothermometer. We summarized daily midpoint temperature (i.e., midpoint between minimum and maximum daily temperatures) and annual mean daily maximum temperature across one annual cycle as an indicator of temperature extremes at each site. These two variables were previously demonstrated to represent separate climatic phenomena at Fongoli (Wessling et al. 2018b). Unfortunately, consistent rainfall data were not collected for a full annual cycle across three of the six sites. We therefore extracted mean annual rainfall (years 1970-2000) from the global BIOCLIM dataset (Fick & Hijmans 2017) at approximately 1 km resolution (30’ latitude), and evaluated these averages relative to the three sites for which we could reliably measure daily rainfall across an entire year (Fongoli, Dindefelo, Kayan) using a rain gauge located at each research station.

### Data Analyses

To evaluate potential habitat differences among sites, we summarized habitat characteristics in several ways. We calculated tree density using the number of all trees located within the vegetation plots divided by total area of vegetation plots surveyed at each site. To contextualize floral assemblages at each site, we discuss tree genera within the context of chimpanzee dietary composition. As Fongoli is the only habituated community within our sample and therefore the only site from which a full catalogue of diet is available (Pruetz 2006), we assumed that the Fongoli diet was representative of the diets of all other communities in our sample. Several lines of evidence support significant dietary overlap across these sites (Wessling et al. 2019; Pan African Programme, unpublished data). We therefore categorized tree genera according to the following potential dietary categorizations: consumed fruit genera, consumed non-fruit genera, non-consumed genera, and the *post-hoc* addition of non-consumed and consumed *Acacia* species. We added the last two categories related to the *Acacia* genus following the in-field observation of significant amounts of *Acacia* trees at Hérémakhono, and therefore divided these categories to evaluate *Acacia* distribution across latitudes. Only two *Acacia* species are consumed by Fongoli chimpanzees for their dry fruits (*A. ehrenbergiana* and *A. polycantha*). As chimpanzees are ripe fruit specialists and arguably prefer fleshy fruits over dry fruits, we also calculated tree density of all fleshy fruit species that fall within the Fongoli diet (Pruetz 2006). We define fleshy fruits as fruits that contain a soft pulp or juice at the time of consumption, although this excludes dry fruits like *Adansonia* which is a preferred food species for the Fongoli chimpanzees (Pruetz 2006). We provide a list of fruit and fleshy fruit genera in the Electronic Supporting Material (ESM).

We used site means of number of trees per vegetation plot as a measure of tree abundance and distribution across each site, and therefore as a proxy of landscape characteristics or heterogeneity of vegetative types within the landscape (Figure 2). If total tree density is constant among the sites but the standard deviation in the number of trees per plot varies, then the distribution of the same number of trees within the landscape will likewise vary. The number of trees per plot can also be used as an objective measure of habitat classification. Habitat classifications are often subjective and can suffer from issues of consistency and inter-study disagreement (van Leeuwen et al. 2020, this issue). We therefore assigned objective thresholds of 50% or 66% or more of the maximum number of trees per plot in the dataset (32 trees) to compare the distribution of potentially closed canopy type habitats among the sites, with the assumption that plots exceeding this threshold are similar to closed canopy habitats like gallery forest. Due to the structure of the Fongoli data (one continuous transect), we did not include Fongoli in these analyses, as the savanna-mosaic ecoregion is markedly heterogeneous in vegetative structure and a transect 3.4 km in length is unlikely to accurately reflect site-level tree density and composition (see ESM). Additionally, subsampling of the transect will not result in a sufficient number of spatially independent plots to accurately estimate these metrics, as transect length would allow for only 34 plots at the Fongoli site and a minimum number of 100 spatially-explicit plots per site is needed to estimate these characteristics in this landscape (see ESM for information on minimum sampling thresholds). While sampling at Kayan (N=93) also did not exceed this threshold, it is still likely to estimate these metrics with moderate precision (see ESM; Table S1, Figure S2). We therefore include Kayan but not Fongoli in these analyses.

**Figure 2.**
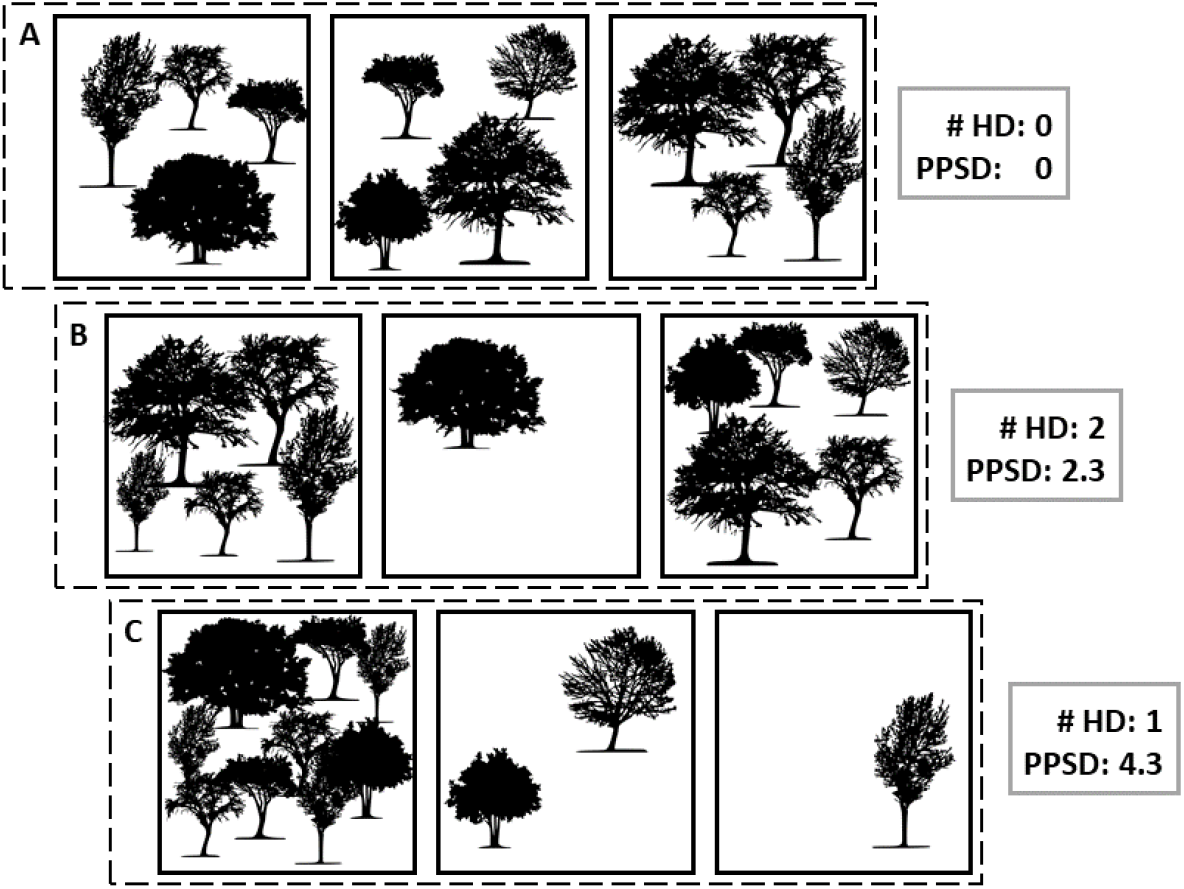
Example of variation in landscape characteristics in three fictional sites with three vegetation plots each. All three sites have identical tree density (12 total trees / area) and mean number of trees per plot (4 trees per plot). The SD of trees per plot varies across sites (PPSD), as an indicator of habitat heterogeneity within a landscape, as does the number of high-density plots (# HD; defined as minimum five trees per plot).

All analyses were conducted in the statistical software R (version 3.6.1; R Core Team 2019). To evaluate potential range limit effects on chimpanzee densities and a number of potentially relevant environmental variables across the six sites, we conducted Spearman’s rank correlation tests with the centered latitude of each site’s grid system as a predictor. These environmental variables included mean and maximum daily temperatures, rainfall, total basal area, total and fleshy fruit tree species density, percentage of high-density plots, number of food and fruit genera, and percentages of food categories. To resolve the issue of dichotomized decisions about significance at a fixed threshold we report p-values between 0.05 and 0.1 as a ‘trend’ for all models (Stoehr 1999).

Our assessment of the Spearman’s rank correlations between the various environmental variables with latitude was hampered by issues of multiple testing of potentially interrelated environmental variables. To tackle this issue we used a combination of Fisher’s omnibus test (Haccou & Meelis, 1994; Quinn & Keough 2002, P. 50) and a permutation test (Adams & Anthony 1996). This method was developed to account for the fact that when researchers wish to evaluate the relationship of one variable with a particular outcome (here, broadly the environment and latitude), they may do so by using a number of potentially non-independent covariates which comprise that outcome (e.g., various ecological and climatic components; Potter & Griffiths 2006).

Following this method, we first determined the exact P-value for each of the correlations (Siegel & Castellan 1988; Mundry & Fischer 1998), and then condensed them into a single quantity for the set using X^2′^ = −2 × Σ(log_e_ *P*) where log is the natural logarithm. If these P-values were independent (i.e., each test’s results independent from all others), we could assess the significance of the resulting quantity by comparison with a chi-square distribution, but the lack of independence of P-values (i.e., potential collinearity among environmental variables) invalidates this approach. We overcame this constraint using a permutation test which shuffles the six possible latitudinal values across all possible combinations of sites while keeping the associations of the environmental variables within sites unaffected (keeping their potential non-independence, e.g., same values of BA and rainfall at ‘Makhana, but ‘Makhana’ randomly assigned any one of the six potential latitudes). The permutations of latitude were exact; that is, we enumerated all 720 possible permutations of the six latitude values. A complication arose from the fact that we did not record six of the environmental variables at Fongoli (tree and fleshy fruit tree density, BA, number of trees per plot, high density plots-66%, high density plots-50%). For the environmental variables comprising the missing value we did not include Fongoli’s latitude in the permutation. We then determined the exact P-value for each of the correlations between the environmental variables and the permuted latitude and combined them into χ^2′^ as described above. We determined the final P-value for the overall association between latitude and the environmental variables as the proportions of permutations revealing a χ^2′^-value at least as large as that of the original data, essentially a measure of the likelihood that a given combination of environmental gradients would accidentally (i.e., assuming no correlation between latitude and the environmental variables) correlate equally or more strongly with latitude than the original configuration. We implemented this test in R with the aid of the function *permutations* of the package ‘gtools’ (version 3.8.1; Warnes et al. 2018).

In all analyses, we used the Universal Transverse Mercator coordinate system (UTM) as a proxy for latitude, as UTM coordinates are scaled in meters and all sites occupied two adjacent UTM zones (28P and 29P) for which northings correspond across zones. To test for potential latitudinal differences in DBH of trees across our sample, we fitted a linear mixed model (LMM: Baayen 2008) with a Gaussian error distribution using the ‘lme4’ package in R (Bates et al. 2015) with genus and site as random effects, and the latitude of each individual tree as a test predictor. We lacked specific location information on several trees at Fongoli (n=264) and elsewhere (n=16) and therefore assigned the latitudinal midpoint of the respective site. As the distribution of the response variable was highly skewed, we considered using a generalized linear mixed model (GLMM) with Gaussian error and log link function, however this model severely violated assumptions about normally distributed residuals. Therefore, we chose the LMM with a log transformed DBH response to meet model assumptions of normally distributed and homogenous error. We initially tested the potential latitudinal effects of number of trees per vegetation plot in a similar manner using a GLMM with Poisson error distribution, with latitude of each vegetation plot as a test predictor and site as a random effect. However, this model suffered from overdispersion (which can lead to increased type I error rates: Gelman & Hill 2007) and complete separation issues. We therefore fitted the model using a Negative Binomial error structure instead, without the random effect of site and found this resolved both issues. To aid comprehension of the model estimates, we z-transformed latitude for both models. We compared the fit of the full models to their respective null models using a likelihood ratio test (Forstmeier & Schielzeth 2011). Each null model was identical to the full model except it lacked the test predictor, latitude. Prior to fitting the DBH model, we checked for deviations from model assumptions of normally distributed and homogenous residuals using visual inspection of qq-plots and residuals plotted against fitted values. We assessed model stability by excluding levels of the random effects one at a time and comparing the estimates derived from these datasets with those derived for the full dataset. We did not identify any issues with both final models. We estimated effect sizes of both models using the function *r.squaredGLMM* of the package ‘MuMIn’ (Barton 2019), and report the variance explained by the fixed effects (marginal R^2^_m_) and the fixed and random effects (conditional R^2^_c_; Nakagawa & Schielzeth 2013). Our dataset for the DBH model included 7200 trees over six sites and 74 genera, whereas data for the vegetation plot model included a total of 873 vegetation plots across five sites.

### Ethical Note

The research presented here was non-invasive and did not directly involve research on any animal subjects. All research, human and non-human, was approved by the Max Planck Society, and permission for this research was granted by the Direction des Eaux, Forêts, Chasses et de la Conservation des Sols in Senegal. The authors declare that they have no conflict of interest.

## RESULTS

The discovery of night nests during reconnaissance surveys (i.e., recces) confirmed chimpanzee presence at all surveyed sites (Table 1). Surveyed transects averaged 58.5 km per site (range: 20-94 km), and we discovered chimpanzee nests on all transects except at Hérémakhono. ESW was 32.80m ± 3.73 (SE), based on the best fit of the half-normal key function with a cosine adjustment of order 2 and 3 to our data (Figure S1). Generally, we discovered far fewer nests at Hérémakhono than at all other sites (Table 1). Across transects in which chimpanzee nests were discovered, chimpanzee densities averaged 0.11 individuals km^−2^(range: 0.05 – 0.29 individuals km^−2^). A sufficient number of nests were encountered at these sites (mean ± SE: 89.6 ± 15.0, range: 66 - 147) to reliably estimate chimpanzee densities (Kühl 2008). Overall, chimpanzee densities declined significantly with increasing latitudes (Table 1).

Overall, we found a statistically significant association between the environmental variables and latitude (Fisher’s omnibus test in combination with permutation procedure: χ^2′^= 75.57, P=0.028), indicating broad ecological difference within our measured variables across these six sites (Figure 3).

**Figure 3.**
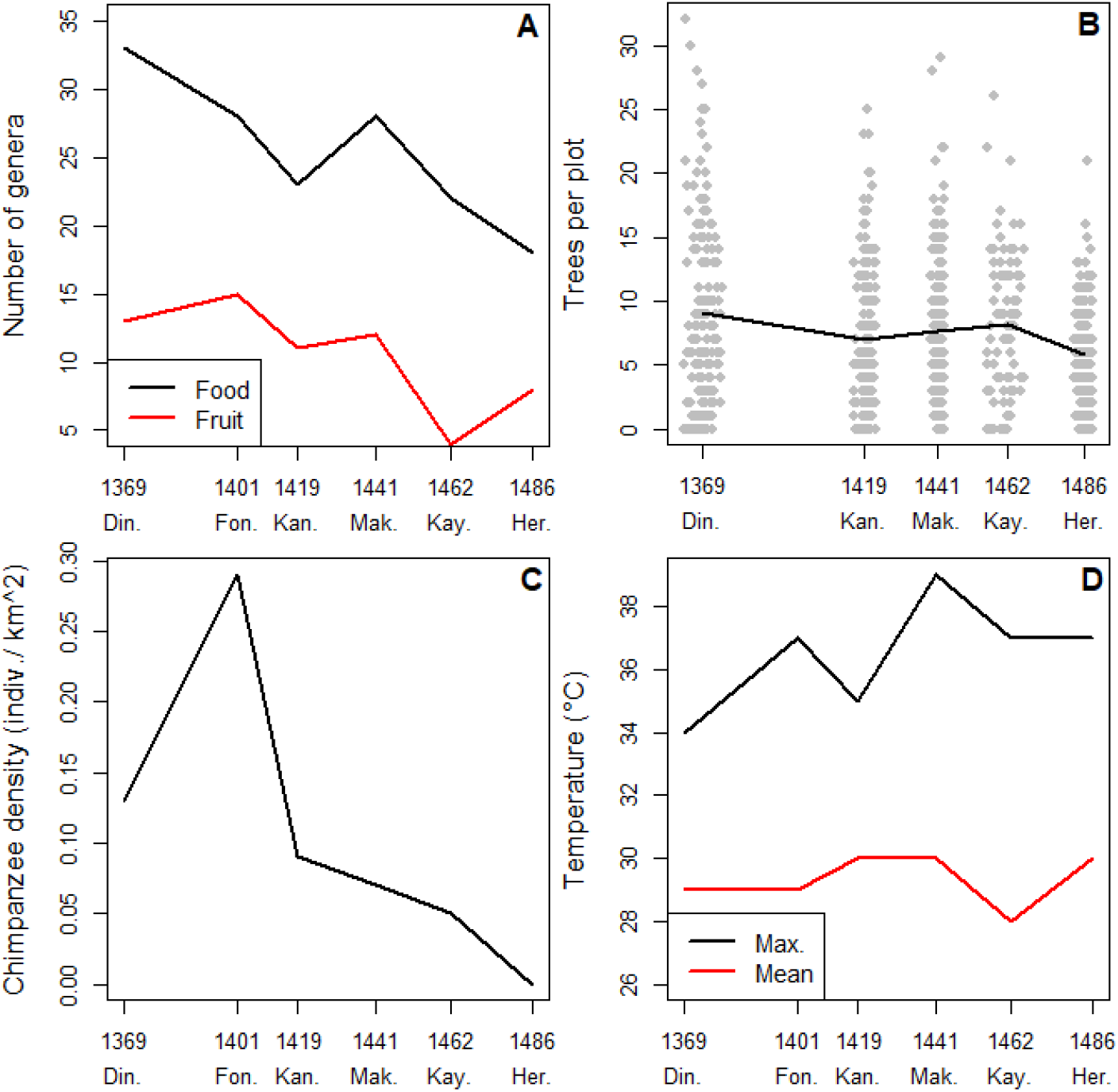
Latitudinal variation in (A) number of genera (B), number of trees per plot, (C) chimpanzee (*Pan trodlodytes verus*) density, and (D) daily temperature across the six study sites (Hérémakhono, Kayan, Makhana, Kanoumering, Fongoli, Dindefelo; 2012-2017). X-axes show latitudinal midpoints for each site (rounded to the nearest 10000 m: UTM Zone 28P), with the exception of datapoints in panel B, which are located at true latitudes of each plot.

**Figure 4.**
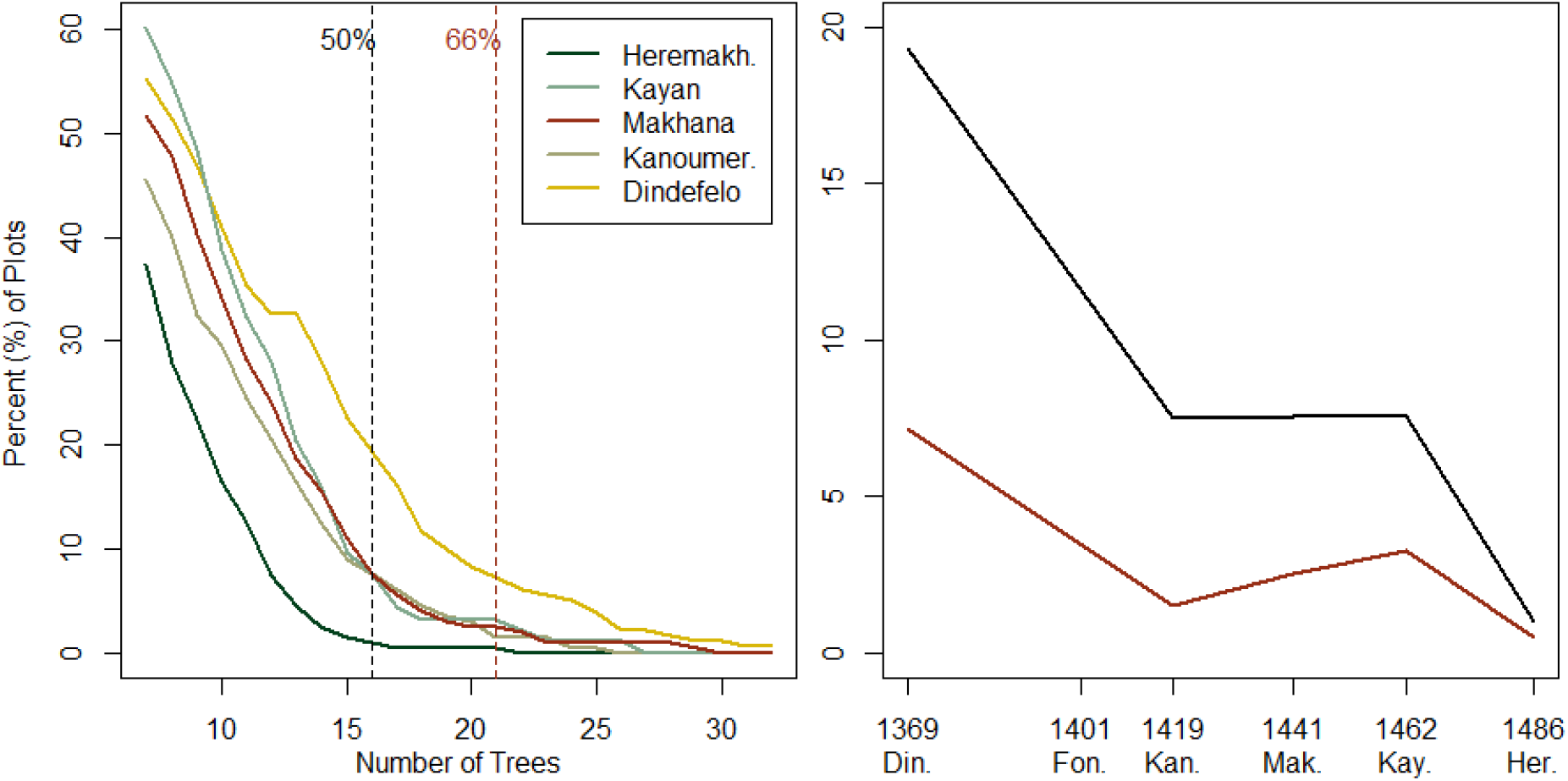
(Left) Percentage of vegetation plots (y-axis) containing a minimum number of trees (x-axis, range from approximate average number of trees per plot (7 trees) to maximum number of trees per plot observed in the dataset (32 trees; 2012-2017). Vertical lines represent 50% and 66% of the maximum number of trees per plot. (Right) Changes in percentage of 50% (black) and 66% (red) high density plots across chimpanzee (*Pan troglodytes verus*) sites (Hérémakhono, Kayan, Makhana, Kanoumering, Fongoli, Dindefelo) and latitudes. X-axis depicts latitudinal midpoints for each site (rounded to the nearest 10000 m: UTM Zone 28P).

The average midpoint temperature across the six sites was 29.3 °C, and the average daily maximum temperature was 37.0 °C (Table 1). Overall, we observed no significant latitudinal patterns in temperature midpoints for daily mean temperature, as well as mean daily maximum temperature (Table 1). Average annual rainfall from global datasets for these sites was 1009 mm ± 133 (SD; range: 849 – 1220mm) and we found a significant latitudinal decrease of over 350 mm in rainfall across all sites (371mm over 126.1km distance; Table 1). Values from global datasets corresponded well to our measurements of rainfall from 2013 for both Dindefelo and Fongoli, but underestimated rainfall measurement by 200 mm at Kayan. Mean measured rainfall for the region was 1123 ± 66 (SE) mm year^−1^(n=3 sites).

We recorded data for 7209 trees over a 41.7 ha sampling area across the six sites (Table 2). Tree density across our dataset averaged 188 trees ha^−1^(range:145 – 226 trees ha^−1^) but showed no consistent statistical pattern in density across latitudes (Table 2). As a measure of preferred food tree species density, fleshy fruit tree densities averaged 45.6 trees ha^−1^across the dataset (range: 25.9 - 66.2 trees ha^−1^), with Hérémakhono containing the lowest density, 30% lower than the density of the site most similar in density. However, we found no consistent statistical pattern of fleshy fruit tree densities across a latitudinal gradient (Table 2).

**Table 2.** Vegetation plot summaries for six chimpanzee (*Pan troglodytes verus*) sites in Senegal (2012-2017), ordered from north (top) to south (bottom). Significant (p<0.05) latitudinal patterns (based on Spearman correlations between the variable and latitude) are indicated in underscored bold text.

| Site | Total number of trees | Number of vegetation plots | Tree density (indiv. ha<sup>-1</sup>) | Fleshy fruit tree density (indiv. ha<sup>-1</sup>) | Average number of trees per plot ± SD | % high density plots (50%*) | % high density plots (66%**) | Average DBH (cm) ± SD*** | Overall BA ha<sup>-1</sup> (m<sup>2</sup> ha<sup>-1</sup>) |
| --- | --- | --- | --- | --- | --- | --- | --- | --- | --- |
| Hérémakhono | 1162 | 201 | 144.5 | 27.4 | <u>**5.77 ± 3.68**</u> | 1.00 | 0.50 | <u>**19.3 ± 11.7**</u> | 6.38 |
| Kayan | 758 | 93 | 203.8 | 48.7 | <u>**8.15 ± 5.41**</u> | 7.53 | 3.23 | <u>**19.2 ± 10.3**</u> | 7.50 |
| Makhana | 1530 | 199 | 192.2 | 53.4 | <u>**7.69 ± 5.49**</u> | 7.54 | 2.51 | <u>**19.3 ± 13.2**</u> | 9.75 |
| Kanoumering | 1398 | 200 | 174.8 | 40.4 | <u>**6.99 ± 5.35**</u> | 7.50 | 1.50 | <u>**19.2 ± 11.0**</u> | 6.94 |
| Fongoli | 727 | - | - | - | - | - | - | <u>**19.2 ± 16.0**</u> | - |
| Dindefelo | 1637 | 181 | 226.1 | 74.4 | <u>**9.04 ± 7.19**</u> | 19.33 | 7.18 | <u>**18.2 ± 10.9**</u> | 9.29 |
| *p* - value |  |  | 0.35 | 0.23 | (see | 0.23 | 0.23 | (see | 0.45 |
| *rho* ($\rho$) | - | - | -0.60 | -0.70 | Table 3) | -0.70 | -0.70 | Table 3) | -0.50 |

\* Corresponds to 50% of the maximum 32 trees per plot (16 or more trees per plot)
\*\* Corresponds to 66% of the maximum 32 trees per plot (21 or more trees per plot)
\*\*\* Site mean controlled for tree genus

Across the dataset, we measured an average of 7.4 trees per plot (range: 0 to 32 trees per plot). Investigating whether the number of trees per plot varied across a latitudinal gradient, we found the number of trees per plot correlated significantly and negatively with latitude, indicating fewer trees per plot as plots increased in latitude (Table 3). This corresponded to a mean decrease in 2.7 trees per plot over the measured latitudinal range (126.1 km), although the explained variance in trees per plot by this model was exceptionally low (R^2^_m_: 0.028, R^2^_c_: 0.018). This pattern appears to be driven by Hérémakhono, which averaged 5.8 trees compared to site means of 7.0 - 9.0 trees per plot at the other sites, while likewise being more consistent in number of trees per plot over the site as a whole, with standard deviation of Hérémakhono plots at 3.7 trees per plot compared to 5.4 - 7.2 trees per plot at the other sites.

Hérémakhono had considerably smaller percentages of high-density habitats (1.0% of plots at 50% of the maximum number of trees; Table 2, Figure 3) relative to the four other sites (range: 7.5-19.3% plots), and this paucity remained consistent regardless of the choice of threshold used to define a high density plot (Figure 3). This pattern, however, was non-significant (Table 2). If we defined high-density plots as 66% of the maximum, Hérémakhono appeared to offer almost no closed-canopy habitats (0.5%), whereas all other sites offered at least small fractions (range: 1.5-7.2% plots). This pattern was also non-significant (Table 2). Despite the lack of statistical significance, both thresholds followed clear decreases from one latitudinal extreme to the other. These results were comparable to on-the-ground observations during the study that Hérémakhono tree distribution appeared relatively even throughout the study area, but with a complete absence of closed-canopy type habitats.

**Table 3.** Model results for the effect of latitude on DBH (cm) and number of trees per plot at six chimpanzee (*Pan troglodytes verus*) sites (Hérémakhono, Kayan, Makhana, Kanoumering, Fongoli, Dindefelo; 2012-2017). Estimates and standard errors are back transformed to their original scales.

| Term | Estimate ± SE | $\chi^2$ | p |
| --- | --- | --- | --- |
| **DBH (cm) across site (LMM; log transformed)** |  |  |  |
| (Intercept) | 18.7 ± 1.0 | - | - |
| Latitude* | 1.0 ± 1.0 | 6.912 | 0.009 |
| **Number of trees per plot (Negative binomial)** |  |  |  |
| (Intercept) | 7.4 ± 1.0 | | |
| Latitude** | -0.9 ± 1.0 | 20.463 | <0.001 |

\* z-transformed, mean ± SD at the original scale: 1425943 ± 39802
\*\* z-transformed, mean ± SD at the original scale: 1433728 ± 40715

We identified 78 unique genera across the dataset, with an average of 39 ± 4 (SE) identified at each site (range: 30 – 58 species). The number of food genera available to chimpanzees decreased significantly with increasing latitude (Table 4). This pattern was likewise significant for fleshy fruit tree genera, which averaged 11 ± 2 (SE) genera across sites (range: 4 – 15 genera). Hérémakhono offered the fewest food and among the fewest fruit species of all sites, nearly half of that in our southern-most site, Dindefelo. The only site which had fewer fruit species was Kayan. We cannot discern whether this pattern originates from under-sampling of plots or true ecological difference (ESM).

**Table 4.** Percentage of trees within each dietary category, and number of genera per chimpanzee (*Pan troglodytes verus*) site (2012-2017). Significant (p<0.05) latitudinal patterns are indicated in underscored bold text.

| Site | Number of consumed genera (# of fleshy fruit genera) | Percentage of trees per dietary category (%) |  |  |  |  |
| --- | --- | --- | --- | --- | --- | --- |
|  |  | Fruit | Non-fruit | Non-consumed | Non-consumed (*Acacia*) | Consumed (*Acacia*) |
| Hérémakhono | <u>**18 (8)**</u> | 31.6 | 43.9 | <u>**13.1**</u> | 1.5 | 10.0 |
| Kayan | <u>**22 (4)**</u> | 39.4 | 40.1 | <u>**12.8**</u> | 0.7 | 7.0 |
| Makhana | <u>**28 (12)**</u> | 38.3 | 45.8 | <u>**14.9**</u> | 0.1 | 0.8 |
| Kanoumering | <u>**23 (11)**</u> | 34.2 | 42.3 | <u>**18.4**</u> | 0.3 | 4.8 |
| Fongoli | <u>**28 (15)**</u> | 42.2 | 29.1 | <u>**28.3**</u> | 0.4 | 0.0 |
| Dindefelo | <u>**33 (13)**</u> | 43.7 | 23.2 | <u>**30.4**</u> | 0.0 | 2.7 |
| *p - value* | <u>**0.015 (0.058)**</u> | 0.103 | 0.136 | <u>**0.017**</u> | 0.103 | 0.136 |
| *rho* | <u>**-0.899 (-0.829)**</u> | -0.771 | 0.714 | <u>**-0.943**</u> | 0.771 | 0.714 |

Of the floristic composition found at each site, an average of 75.6 ± 2.5% (SE, range: 66.9-84.1%) of trees produced at least one plant part known to be consumed by Fongoli chimpanzees. We observed clear increases in number of *Acacia* trees at Hérémakhono relative to other sites, with over 11% of identified trees falling within this genus, relative to a non-Hérémakhono average of 3.4 ± 1.3% (SE; Table 4). An average of 38.2 ± 1.9% (SE) non-*Acacia* trees in our dataset are eaten for their fruits (both fleshy and non-fleshy; range: 31.6 to 43.7%). Hérémakhono had the lowest percent of non-*Acacia* fruit trees (31.6%) but the highest percentage of edible *Acacia* trees (10.0%) in comparison with edible *Acacia* percentages between 0.8 – 7.0% at other sites. We observed a significant decrease in non-consumed species with increasing latitudes (Table 4).

We observed a significant latitudinal effect on tree size, as measured by DBH (R^2^_m_: 0.002, R^2^_c_: 0.453; Table 3). The fitted model estimated a 3.3 cm increase in DBH over the measured latitudinal range (126.1 km). Average DBH across all trees was 18.7 cm (total range: 10 to 250 cm) when accounting statistically for the confounding effect of genus and site, although site averages varied little (range: 18.2 – 19.3 cm; Table 2) and intra-site variation was fairly consistent (SD, range: 10.3-16.0 cm). Overall BA averaged 8.0 ± 0.7 m^2^ha^−1^(SE) across the dataset (range: 6.4 – 9.7 m^2^ha^−1^) but did not follow a significant latitudinal pattern across the sites (Table 2).

## DISCUSSION

We describe here the habitat characteristics of multiple previously undescribed sites in a savanna-mosaic ecoregion and relate this to the structure of the range edge of the western chimpanzee. As predicted, we found that chimpanzee densities declined with increasing proximity to the range limit, and that several habitat characteristics likewise declined in parallel. We observed distinct differences in these measures in particular in the northernmost site (Hérémakhono) from the more southern sites, providing additional support that Hérémakhono likely represents the last vestige of chimpanzee occupation at the limit. These insights have the potential to further inform us as to the structure of the chimpanzee distributional limit and the potential limits to chimpanzee niche tolerance overall.

The habitat characteristics of the savanna-mosaics described here offer a point of comparison to results from chimpanzees living in more forested habitats (Potts et al. 2009; Bortolomiol et al. 2014; Potts and Lwanga 2013). In comparison to average tree size (as measured in DBH) in forested habitats (e.g., feeding trees: Chapman et al. 1995; Tweheyo & Lye 2003; Janmaat et al. 2016), tree size appears to be overall smaller in our savanna-mosaic dataset. Additionally, basal area coverage of our dataset confirm that these landscapes harbor less tree coverage compared to forested sites (Potts and Lwanga 2013; Bortolomiol et al. 2014) as is expected based on global patterns (Crowther et al. 2015). As such, the assumption that these habitats offer reduced food availability than forested habitats is broadly confirmed if measured by basal area alone; however, direct phenological comparison suggests tree abundance may not reflect the best measures of food availability as food production rates may vary across landscapes (Wessling et al. 2018a). Additionally, savanna-mosaic landscapes appear to offer on average fewer genera (average 39 sampled genera per site; this study) than forested habitats (e.g., 66 genera: Potts and Lwanga 2013), underlining why chimpanzees living in savannas have a narrower diet (Pruetz 2006; Webster et al. 2014) than those of their forest-dwelling counterparts (Watts et al. 2012; Wrangham 1977).

We observed a pattern of chimpanzee density decline over approximately 126 km within a single ecoregion, suggesting that chimpanzee biogeography may conform to abundant center niche patterns (Sexton et al. 2009), with highest chimpanzee densities towards the center of their range. Higher density estimates from sites farther south in Guinea and Guinea-Bissau (e.g., Sousa et al. 2011; WCF 2016; Kühl et al. 2017) further extend this gradient within the biogeographical range of the subspecies. Environmental conditions broadly varied with decreasing distance to the chimpanzee range limit, and a few potential contributors of environmental drivers of this limit lie amongst our metrics (e.g., reduced food species diversity, refuge from heat, and water availability). One argument is that chimpanzees in this region require diverse plant species communities to support their diverse diet (Kortlandt 1983); such a hypothesis is supported by evidence from East Africa demonstrating that fruit species richness (Balcomb et al. 2000), especially fruit species that produce fruits during periods of low food availability (Potts et al. 2009; Bortolamiol et al. 2014), are predictors of chimpanzee densities. Fruit species diversity, specifically fleshy fruit tree diversity, may therefore be a limiting factor across the chimpanzee range in not only determining local chimpanzee densities (Balcomb et al. 2000; Potts et al. 2009; Bortolamiol et al. 2014), but also determining their biogeographical limits. We find initial evidence that such a pattern may hold in this savanna ecoregion, as floristic diversity declined with decreasing distance to the range limit.

The mechanism by which food species richness may influence chimpanzee distribution is in the restriction of the number of food choices available, especially when preferred food items become scarce. Chimpanzees switch to less preferred food items (i.e., non-fruit items) when preferred food items are not available (Wrangham et al., 1991; Furuichi et al., 2001). If these food items are also constrained, they may need to switch to fallback foods. For example, Webster et al. (2014) concluded that reduction in dietary diversity likely drove the Toro-Semliki chimpanzees to higher rates of insectivory. If chimpanzees at Hérémakhono have fewer overall food and fleshy fruit genera available to them, they are likely to face more frequent or pronounced periods of food scarcity. The increases in food patch size (i.e., increased DBH) we observed may help to offset these constraints at Hérémakhono, but patch size can only offset abundance constraint as long as ephemeral food patches remain continuously present.

Hérémakhono chimpanzees do appear to avoid severe nutritional deficits (as evidenced by δ^15^N comparisons) by depending on fallback foods exceptional to dietary patterns of other chimpanzees in the region (^13^C enriched dietary items e.g., C_4_ grasses or domestic crops: Wessling et al. 2019). One potential candidate may be an increased reliance on *Acacia* food items, as the Hérémakhono flora was disproportionately comprised of consumed *Acacia* species relative to the other sites. Although *Acacia* trees are important food items for some savanna primates (Barnes 2001; Isbell et al. 2013) they are infrequently consumed at Fongoli (Pruetz 2006). That Hérémakhono chimpanzee density was so low suggests that *Acacia* is likely to be an insufficient fallback food to compensate for restricted food species diversity in this landscape and that these chimpanzees may already be stretched to the edge of their dietary flexibility.

While intra-annual variation in fruit availability is a frequent consideration in explaining chimpanzee behavior and physiology (e.g., Chapman et al. 1995; Boesch 1996; Murray et al. 2006; Wittiger & Boesch 2013; Samuni et al. 2018; Wessling et al. 2018a,b) it is an underappreciated predictor of chimpanzee distribution across sites, despite clear indications that it is an important determinant of other frugivorous primate distribution (e.g., Marshall et al. 2009b, 2014). Our results suggest that quantification of fruit species assemblages comprise an important measure of food availability consistency and dietary tradeoffs as potential limitations to chimpanzee distribution. In this sense, chimpanzee would appear to be subject to the same constraint generally considered to limit primate distribution—food availability during periods of food scarcity (e.g., Potts et al. 2009, Marshall & Leighton 2006; Marshall et al. 2009a).

In an extreme ecoregion for chimpanzees like the savanna-mosaics of southeastern Senegal, potential dietary limitations of floral assemblages form only part of the picture. In these landscapes, closed canopy habitats are rare and frequently fall below 5% of total land coverage, with preferred habitats like gallery forest covering closer to 2% (McGrew et al., 1988; Bogart & Pruetz 2008; Pruetz & Bertolani 2009; Lindshield et al. 2019). These habitat types are especially important, as Fongoli chimpanzees preferentially spend their time in these habitats, presumably as a means of behavioral thermoregulation (Pruetz & Bertolani 2009). If our estimation of high-density plots serves as a suitable proxy for these habitat types, then these habitats are consistently relatively rare across our sites, with a drop-off at the northern reaches. As such, it is likely that landscapes at the northern limit offer fewer refuges for chimpanzees to avoid the high temperatures (37 ° C average daily maxima) than those further to the south. Although at a site level tree density at Hérémakhono was similar to that of the other sites, average number of trees per plot was lowest and least variable, with the percentage of high-density plots likewise close to zero. Hérémakhono therefore was rather uniform in habitat types available and offered little to no shaded habitats for chimpanzees under the same thermal challenges as those further from the range limit, an observation we also made during data collection. As thermoregulatory stress is particularly constraining for chimpanzees in Senegal (Wessling et al. 2018b), we find some support for the hypothesis that the lack of thermal refuge contributes to the chimpanzee range limit (McGrew et al. 1981), especially as obligatory resting time due to thermal constraints is thought to be a limit to primate biogeography (Korstjens et al. 2010). In this sense, minimum number of resting opportunities (i.e., refuge locations) may be another regulating component of chimpanzee (and likely other species’) biogeography.

Water availability has also been identified as a general factor dictating chimpanzee site suitability in Senegal (Lindshield 2014), and any reduction in water availability is likely to directly exacerbate an already significant constraint to chimpanzees in this landscape (Wessling et al. 2018b). Our results indicated significant decreases in average rainfall patterns with increasing latitude, matching the habitat differences. Floristic assemblages in tropical ecosystems are strongly influenced by rainfall patterns (Bongers et al. 1999; Engelbrecht et al. 2007), and the increase in arid-adapted arboreal flora like *Acacia* species likewise corroborate evidence of increasingly arid conditions at the northern limit of the chimpanzee range. Rainfall therefore appears to play a considerable role in shaping mechanistic limitations to savanna chimpanzee distribution, in that it directly influences water availability to chimpanzees in a thermally challenging environment, while likewise shaping floristic (and therefore potential dietary) composition.

It is possible that all three of the constraints we have discussed (floral assembly/dietary availability, opportunities for thermal refuge, and water availability) contribute collectively as proximate determinants of the chimpanzee range limit. While we can consider the ecological variation described here within the context of longer-term, coarse-grained patterns previously described at national and continent-wide scales (Simpson 1964; Rosenzweig 1995; Crowther et al. 2015), all of our sites are outside formal protection zones and experience some degree of anthropogenic disturbance. Although the Shield ecoregion remained relatively stable in vegetation cover relative to other ecoregions (Tappan et al. 2004), it is unclear if floristic communities are still regionally stable. Nomadic pastoralists in the region specifically target tree species key to chimpanzees (Massa 2011) and felling for livestock fodder was evident at all six sites. How these and other anthropogenic influences shape the biotic communities chimpanzees inhabit and how this varies across the range edge will inform us on the role humans play in dictating chimpanzee distribution locally and regionally. Regardless of the origin of the ecological patterns described here, both natural and anthropogenic pressures lead to the same pattern of environmental drivers of chimpanzee decline at the range limit. Additional investigation into patterns of direct anthropogenic or predatory influence on chimpanzee density at the range edge will likewise be informative as to the relative importance of direct top-down processes in comparison with bottom-up environmental correlates.

Chimpanzee densities appear to be sustained close to the range edge with apparent rapid declines at the limit. Complementary evidence suggests that the habitat of Hérémakhono differs significantly enough in its biotic structure that it may potentially fail to support a full chimpanzee community. The exceptionally low chimpanzee encounter rates and the fact that no nest groups larger than three nests (Wessling, unpublished data) were discovered at Hérémakhono indicate that the chimpanzees living at this site form an exceptionally small social unit. While we do not have data to indicate if this landscape represents a demographic sink, it is nonetheless likely that Hérémakhono represents a form of distribution ‘bleed-over’ and may serve as the marginal transition zone between habitats for which chimpanzees are adapted (e.g., Fongoli: Wessling et al. 2018b) and those in which they are not. These habitats may be temporarily attractive to migrating individuals, for example, as a means to reduce competition, and therefore may even be examples of ecological traps (Battin 2004). Although we do not have demographic data to investigate whether these habitats negatively impact chimpanzee reproductive success and survival, our results indicate that the habitat of Hérémakhono may represent a population sink. Further investigation into the permanence, behavioral ecology, demography, and movement of chimpanzees around these locations may better inform us on the population dynamics of the range edge.

If additional evidence supports Hérémakhono as a marginal habitat or transition zone (Kawecki 2008), the Kayan region would represent the limit of the chimpanzee fundamental niche, although it is not the biogeographic limit of chimpanzee distribution. Such biogeographic ‘bleed-over’ has significant implications for species’ ENMs and predictive models that use species’ distribution patterns to estimate potential suitable habitat in areas where occurrence is unknown. Data from bleed-over regions, or population sinks more generally (Pulliam 1988), are likely to lead ENM models to predict a larger range of suitable environmental conditions than what is sustainable, thereby overestimating suitable habitat coverage for that species.

Our analyses offer exploratory insights into intermediary landscape-level factors between regional level analyses and site-based investigations of chimpanzee habitat characteristics. Regional level analyses often fail to account for smaller-scale processes and variation that may also impact habitat suitability (Abwe et al. 2019), and therefore overlook smaller-scale environmental processes like those we describe here. Our results offer a method of ground-truthing the conclusions of larger-scaled studies and broad-scaled ENMs with regard to the governing environmental variables to chimpanzee distribution. We advocate that similar analyses be conducted to evaluate these patterns at the species level once larger datasets become available. Although species range distributions are often abiotically limited (Pearson & Dawson 2003), we offer several proximate mechanisms through which these limits might be intermediated by biotic patterns for a large bodied organism. We describe several climatically-driven latitudinal patterns on biotic components of the environment (e.g., conversion of floristic composition to arid adapted flora) and biotic contributions to climatic constraints (e.g., refuge from heat via vegetative cover), thereby highlighting additional complexities to range limits likely to be overlooked by broad-scaled ENMs. These complexities suggest that distributional modelling which can integrate data with locally scaled mechanistic implications (e.g., Marshall et al. 2014; Foerster et al. 2016) may be most effective at accurately estimating nuanced distributional constraints for many species, including the chimpanzee.

The approach we use to describe the range dynamics of the chimpanzee may be applied across other species. Analyses like those we present here should allow contextualization of species’ niche patterns over both broader spatial and temporal extents and would ideally allow additional evaluations of the effect of both bottom-up and top-down considerations upon species’ niche limitations simultaneously. Furthermore, the identification of processes dictating species limits as well as patterns explaining distribution or abundance will become increasingly informative in the face of widespread species declines and forecasting the effects of climate change on primates or other species (Martinez-Meyer 2005), as processes of range limitations will dictate a species’ ability to adapt to a changing environment.

