## Supplementary text, tables, and figures for "Chimpanzee (*Pan troglodytes verus*) density and environmental gradients at their biogeographical range edge"

**Electronic Supplementary Material**

**1. Detection Probability of Nests from the Transect**

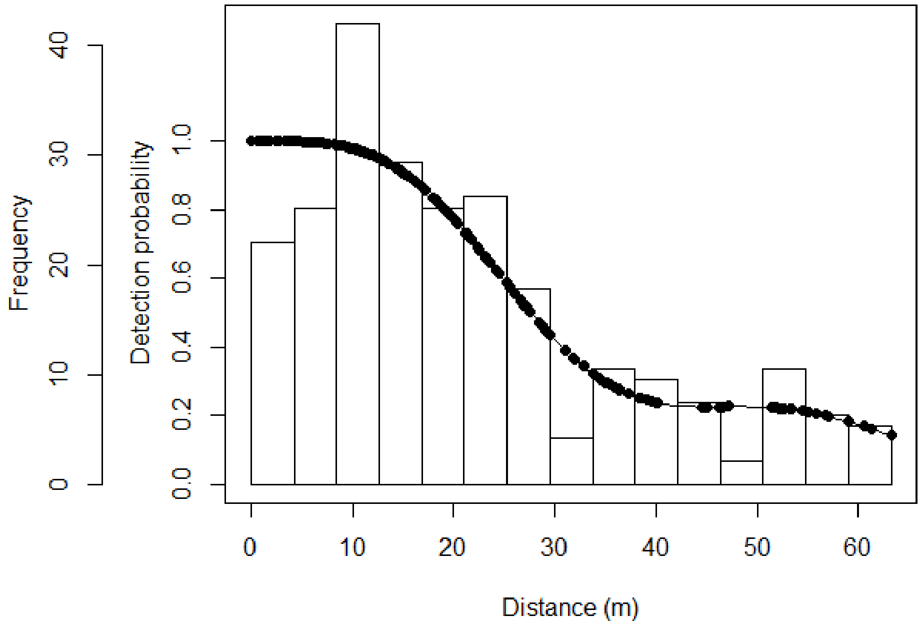


**Figure S1.** Detection probability of half-normal key function with a cosine adjustment of order 2 and 3 model fit (and frequency) to nests encountered along Kanoumering, Makhana, and Fongoli transects.

**2. Genera of Fruit Categories**

Fleshy Fruits: *Lannea, Cordyla, Hexalobus, Sclerocarya, Bombax* (consumed unripe)*, Strychnos, Ficus, Vitellaria, Diospyros, Nauclea, Ximenia, Saba, Quassia, Annona, Spondias*

All Fruits: *Acacia, Adansonia, Afzelia, Allophyllus, Annona, Bombax, Borassus, Ceiba, Cissus, Cola, Cordia, Cordyla, Daniellia, Detarium, Diospyros, Ficus, Grewia, Hexalobus, Landolphia, Lannea, Khaya, Nauclea, Oncoba, Pachystela, Parinari, Parkia, Piliostigma, Quassia, Saba, Sclerocarya, Spondias, Strychnos, Tamarindus, Vigna, Vitellaria, Vitex, Ximenia, Zahna, Zizyphus*

**3. Minimum sampling thresholds for landscape characterization in a savanna-woodland ecoregion**

To ensure that sampling of vegetation plots at our sites was of sufficient effort, we aimed to identify a threshold after which quantification of landscape variables could be considered accurate and representative of true values. As with any metric, under-sampling can lead to errors in precision and accuracy, an error which is likely to be exacerbated in particularly heterogenous datasets (or habitats like savanna-mosaics) or when features of a dataset (e.g. high density plots) are rare. We therefore used the datasets from our four most heavily sampled sites (Dindefelo, Kanoumering, Makhana, and Hérémakhono) to assess a minimum sampling threshold for a reliable estimation of four landscape variables.

To do so, for each number of vegetation plots from 1 to 200 in each site, we simulated 1,000 randomized selections of vegetation plots at each site (i.e., resulting in 1000 x 200 randomized datasets for each of the four sites), with replacement. Then, for each of the randomized datasets we calculated the i) tree density, ii) SD of number of trees per plot, iii) % of high density plots (50%), and iv) number of tree genera per site. The values of these parameters are expected to be the most accurate for 1,000 random selections of 200 vegetation plots, and the least accurate for 1,000 random selections of one vegetation plot.

Following this logic, we used the range of values of the highest randomized sampling effort (i.e., 200 plots with replacement) to estimate a minimum threshold for the number of plots needed for reliable assessment of these habitat variables. For every randomized dataset we quantified the percentage of values that appear outside of the acceptable range (defined as the minimum and maximum of the 200 plot randomized dataset) and set the minimum sampling threshold of vegetation plots when at least 95% of randomized sample sets estimated a value within the acceptable range consistently for every sample size above this threshold. Lastly, to quantify the number of genera per site according to sampling size, we used the ‘specaccum’ function from the *vegan* package (version 2.5-6: Oksanen et al. 2019) using the “random” method and set it to 1000 iterations.

<u>Results and Interpretation</u>

Minimum number of plots required to reach 95% agreement with the plot maximum dataset averaged 82 ± 15 (SD) across all variables but ranged from 45 to 100. For all four sites in our simulations, the number of genera at each site did not reach clear plateaus even by the 200-plot sample size, although this was especially pronounced at sites containing greater number of genera (e.g. Makhana and Dindefelo). These results indicate that 200-plot datasets who are especially diverse are more likely to underestimate true species densities, whereas the extent of underestimation in comparatively less diverse samples (e.g., Hérémakhono, Kanoumering) is shallower. As a result, we are likely to have underestimated latitudinal declines in food species richness we observed in our analyses.

We consider the highest minimum sampling threshold across all variables and sites to be the necessary minimum of vegetation plots needed for our dataset, and therefore recommend that a minimum of 100 vegetation plots (threshold maximum) need to be surveyed in this ecoregion to obtain reliable estimations of landscape characteristics. However, if efforts are especially constrained, a minimum of 82 plots (threshold average) are likely to approach accurate estimates with moderate precision for most sites and variables, but fewer than these number of plots should be considered unacceptable. It is important to note, however, that genus richness is likely to be underestimated by an average of at least 13% (range: 10-16% of the maximum sampled value) even when the 100-plot recommended threshold is reached, as well as also likely to be underestimated still at 200 plots in this ecoregion. If future studies wish to census species richness in these landscapes, it is advisable that they sample greater than 200 plots or use an approach which also targets habitats likely to contain rarer species.

**Table S1.** Minimum number of plots needed to be sampled for 95% of the iterations to fall within the accepted range of the maximum modeled sample (n=200), by site and by environmental metric.

| Site | *Number of plots* | Tree density (indiv/ha) | Percent of high density plots | SD of trees per plot |
|---|---|---|---|---|
| Hérémakhono | *201* | 70 | 45 | 75 |
| Makhana | *199* | 100 | 89 | 77 |
| Kanoumering | *200* | 97 | 75 | 87 |
| Dindefelo | *181* | 83 | 97 | 84 |

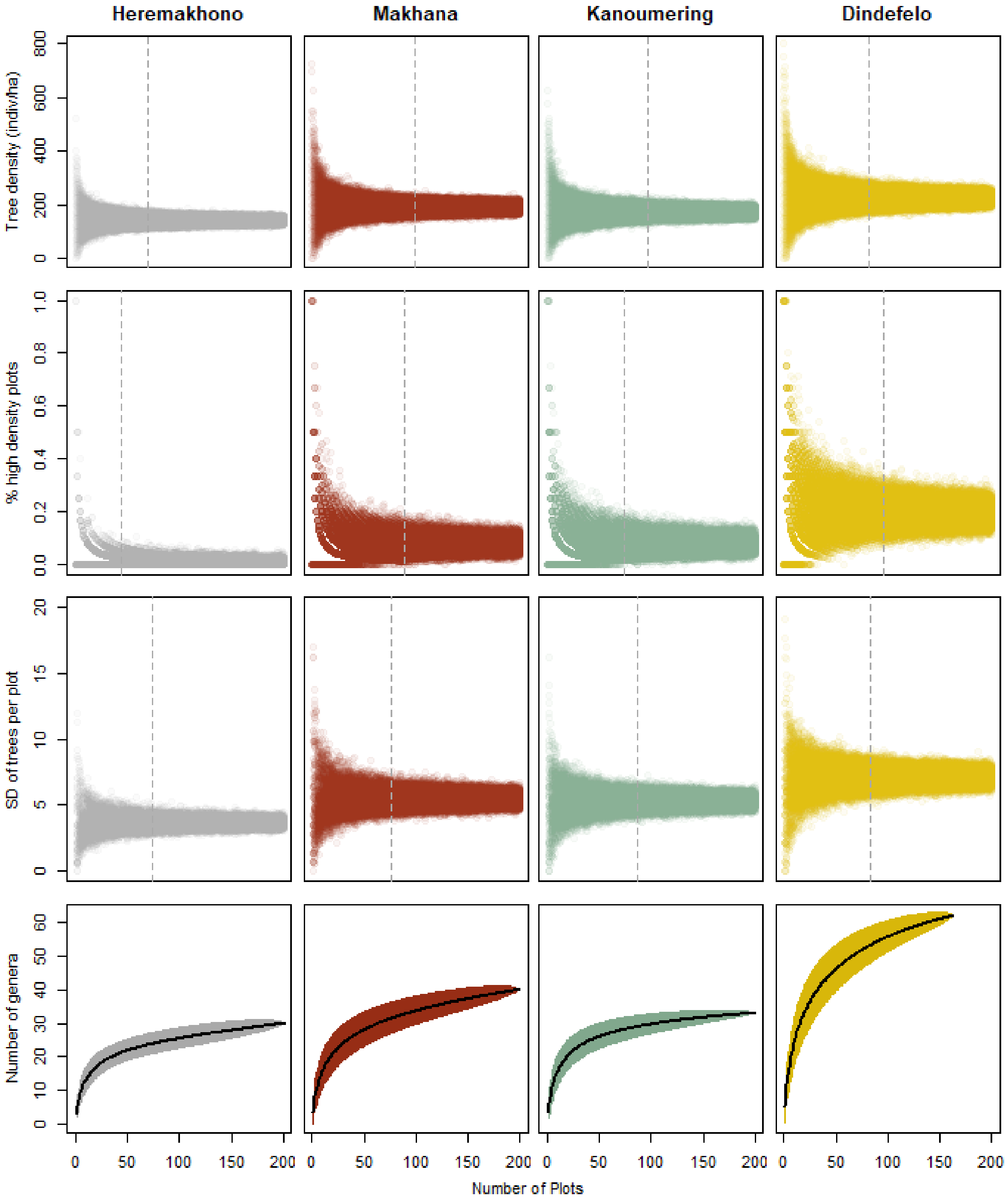


**Figure S2.** Estimates of 1000 iterations per sample size of four environmental variables (tree density, percent high density plots, SD of number of trees per plot, and number of genera) for four datasets. Dotted vertical lines indicate minimum threshold for 95% precision of "true" estimates, black model lines represent sampling averages.
